# Robotically-induced auditory-verbal hallucinations: combining self-monitoring and strong perceptual priors

**DOI:** 10.1101/2022.12.22.521663

**Authors:** Pavo Orepic, Fosco Bernasconi, Melissa Faggella, Nathan Faivre, Olaf Blanke

## Abstract

Inducing hallucinations under controlled experimental conditions in non-hallucinating individuals represents a novel research avenue oriented towards understanding complex hallucinatory phenomena, avoiding confounds observed in patients. Auditory-verbal hallucinations (AVH) are one of the most common and distressing psychotic symptoms, whose etiology remains largely unknown. Two prominent accounts portray AVH either as a deficit in auditory-verbal self-monitoring, or as a result of overly strong perceptual priors. In order to test both theoretical models and evaluate their potential integration, we developed a robotic procedure able to induce self-monitoring perturbations (consisting of sensorimotor conflicts between poking movements and corresponding tactile feedback) and a perceptual prior associated with otherness sensations (i.e., feeling the presence of a non-existing another person). Here, in two independent studies, we show that this robotic procedure led to AVH-like phenomena in healthy individuals, which was further associated with delusional ideation. Specifically, a condition with stronger sensorimotor conflicts induced more AVH-like sensations (self-monitoring), while, in the otherness-related experimental condition, there were more AVH-like sensations when participants were detecting other-voice stimuli, compared to detecting self-voice stimuli (strong-priors). By demonstrating an experimental procedure able to induce AVH-like sensations in non-hallucinating individuals, we shed new light on AVH phenomenology, thereby integrating self-monitoring and strong-priors accounts.

## Introduction

Hallucinations are aberrant perceptual experiences that are reported in several major psychiatric and neurological conditions (Eversfield and Orton, 2019; Insel, 2010; Toh et al., 2015; Waters and Fernyhough, 2017). Hallucinations are of major negative impact. Associated with delusions and psychosis, they often require repeated hospitalization. In neurodegenerative disease, they increase the likelihood of earlier home placement (i.e. Aarsland et al., 2000) and have been linked to higher mortality (Gonzalez et al., 2022). Hallucinations are very common and even occur in 5-10% of the general population, without any medical diagnosis (Larøi et al., 2019). Despite this clinical relevance, hallucination research, and the understanding of the underling brain mechanisms, has been hampered by methodological shortcomings to investigate hallucinations in real-time under controlled experimental settings. A major shortcoming is the absence of controlled procedures allowing to induce hallucinations in laboratory and/or clinical settings (Bernasconi et al., 2022; Pearson et al., 2016). Moreover, hallucination research has predominantly been conducted in clinical populations, and is therefore confounded by comorbidities existing in the tested populations. Recent research has induced hallucinations in controlled laboratory in healthy individuals, using different pharmacological approaches (McClure-Begley and Roth, 2022; Vollenweider and Preller, 2020) and several other procedures (i.e., Flicker-induced phosphenes (Allefeld et al., 2011; Pearson et al., 2016), Ganzfeld effect (Wackermann et al., 2008), sensory deprivation (Mason and Brady, 2009; Merabet et al., 2004), Pavlovian conditioning (A R Powers et al., 2017)).

Hallucinations induced by these methods, however, often have poor ecological validity, and, despite being induced under laboratory conditions and in healthy subjects, are characterized by low experimental control over their content (often not specific), timing and duration (may be long-lasting), and associated with impairments of consciousness (i.e., pharmacological approaches).

We recently described a robotic procedure able to repeatedly induce a specific, highly-controllable and clinically-relevant sensorimotor hallucination – presence hallucination – the sensation that someone is nearby when no-one is actually present, and cannot be seen or heard (Bernasconi et al., 2022, 2021; Blanke et al., 2014). Merging techniques from engineering and neuroscience, this procedure has been used to induce and understand clinically-relevant hallucinations (Bernasconi et al., 2022) under controlled experimental conditions in healthy individuals (Blanke et al., 2014; Dhanis et al., 2022; Orepic et al., 2022, 2021; Serino et al., 2021) as well as in patients with Parkinson’s disease (Bernasconi et al., 2021) and early psychosis patients (Salomon et al., 2020). This method, when combined with auditory-verbal tasks that are carried out during robotic stimulation, is able to induce deficits in auditory-verbal self-monitoring in psychosis patients (Salomon et al., 2020) and induce changes in voice perception in healthy participants (Orepic et al., 2021). Here we asked whether the robotic procedure can be adapted to induce a hallucinatory state in healthy participants that is of major clinical relevance in psychiatry and comparable to auditory-verbal hallucinations (AVH).

AVH, the sensation of hearing voices without any speaker present (commonly known as “hearing voices”) are one of the most common (Bauer et al., 2011) and most distressing (Harkavy-Friedman et al., 2003) symptoms in schizophrenia spectrum disorder. AVH are characterized by a very heterogeneous phenomenology (e.g., varying with respect to voice numerosity, gender, frequency, emotional affect, etc.) (McCarthy-Jones et al., 2014; Woods et al., 2015), and have also been observed in non-help-seeking individuals (Albert R. Powers et al., 2017; Sommer et al., 2010), rendering their diagnosis and treatment challenging. With contemporary treatments being effective only to a certain degree (Lehman et al., 2004), there is a strong need for a better understanding of the underlying mechanisms. Despite the increasing frequency and the large number of studies revolving around AVH, their etiology remains debated, with two prominent and seemingly opposing theoretical accounts, suggesting that AVH result either from 1) deficits in self-monitoring, or 2) overly strong perceptual priors. Although both of these carry some empirical support, only theoretical speculations (Leptourgos and Corlett, 2020; Northoff and Qin, 2011; Swiney and Sousa, 2014; Synofzik et al., 2008; Wilkinson, 2014; Yttri et al., 2022) have been made on how they might coexist in the brain and relate phenomenologically.

The self-monitoring account suggests that AVH arise from a deficit in self-monitoring, more specifically the inability to distinguish self-from other-related events. According to this framework (Miall and Wolpert, 1996; Shadmehr et al., 2010; Wolpert et al., 1995), self-other distinction is achieved by creating sensory predictions related to one’s actions and by comparing them with the actual sensory feedback following those actions. When congruent with the sensory prediction, ascending sensory events are attenuated, and the action is attributed to the self, whereas if the prediction and the ascending sensory events are incongruent, attenuation is reduced and the action is attributed to an external agent (Heinks-Maldonado et al., 2005; Shergill et al., 2003). Impairments in self-monitoring have been observed in schizophrenia (Blakemore et al., 2000; Shergill et al., 2014, 2005), and have been related to psychotic symptoms characterized by a misattribution of self-generated actions (Blakemore et al., 2000; Frith, 1987; Graham-Schmidt et al., 2016), including AVH, explaining them as a misattribution of self-generated speech towards external agents (Feinberg, 1978; Ford et al., 2007; Frith, 1992; Gould, 1948; Green and Kinsbourne, 1990; Green and Preston, 1981; McGuigan, 1966; Moseley et al., 2013). However, the evidence supporting the self-monitoring account for AVH remains largely implicit, as AVH are rarely related to direct experimental manipulations of motor signals and their corresponding sensory feedback. Its empirical support mainly consists of studies (reviewed by (Whitford, 2019)) in which patients with schizophrenia either exhibited a reduced attenuation of auditory neural evoked response while speaking compared to passively hearing their voice, or there was a dysfunctional interregional communication within the speech network (Ford et al., 2012, 2002; Friston and Frith, 1995; Hoffman et al., 2011), both of which were hypothesized to facilitate erroneous feedforward signaling. More direct evidence for the self-monitoring account would consist of a study demonstrating that experimentally-induced self-monitoring perturbations of different degrees (i.e., stronger, weaker) can explicitly lead to different degrees of AVH (i.e., more, less).

The second major account suggests that AVH might be engendered by overly strong beliefs (i.e., priors) about the environment (Cassidy et al., 2018; Corlett et al., 2019; A R Powers et al., 2017; Zarkali et al., 2019). It relies on the predictive coding framework that sees the brain as a hierarchical Bayesian system in which priors (at higher levels) and incoming sensory information (lower levels) are combined for perception (Friston, 2009, 2008). Crucially, precision-weighting of bottom-up (sensory) and top-down (priors) components modulates perception, whereby the component with higher precision dominates perception. Accordingly, hallucinations have been hypothesized to arise when priors carry undue precision, overruling the actual sensory evidence (Adams et al., 2015; Corlett et al., 2019; Fletcher and Frith, 2009; Sterzer et al., 2018). This view is supported by empirical data demonstrating that both clinical (Kot and Serper, 2002) and non-clinical (Alderson-Day et al., 2017) voice-hearers, as well as psychosis-prone individuals (Teufel et al., 2015) favor prior knowledge over sensory information during perceptual inference. These data suggest that perceptual inference in hallucinating individuals is driven by prior beliefs. However, this work has not addressed the relationship of prior beliefs to the perception of voices specifically – i.e., it remains unclear which kinds of priors need to be over-weighted in order to experience AVH, a voice of an external origin. For instance, more direct evidence for the strong-priors account in AVH would consist of a study that experimentally induces a specific perceptual feature (e.g., sensations of otherness in the form of a presence hallucination) and observed a corresponding perceptual bias in AVH (e.g., increased AVH attributed to others).

Here, we developed a new method of inducing AVH in a controlled laboratory environment by integrating methods from voice perception with sensorimotor stimulation, allowing us to investigate the contribution of both major AVH accounts. We used a robotic procedure that can create impairments in self-monitoring as well as elicit the sensations that there is another (alien) person close by (i.e., presence hallucination, PH) (Bernasconi et al., 2021; Blanke et al., 2014; Salomon et al., 2020; Serino et al., 2021). Specifically, we designed a robotic setup that exposes participants to sensorimotor conflicts of various degree between repeated upper-limb poking movements and the corresponding tactile sensations on the back (Hara et al., 2011), linked to the misperception of the source and identity of sensorimotor signals of one’s own body (Bernasconi et al., 2022, 2021; Blanke et al., 2014; Salomon et al., 2020; Serino et al., 2021). Thus, our robotic setup applies sensorimotor conflicts of various degree (i.e., self-monitoring perturbations) and induces a perceptual prior about the presence of an external non-existing agent. We combined this procedure with a voice detection paradigm and measured experimentally-induced AVH-like sensations as an increase in vocal false alarms. Namely, a false alarm in a voice detection task indicates that participants have heard a non-existing voice, rendering vocal false alarms a suitable proxy for lab-induced AVH, as was similarly done by others (Barkus et al., 2011; Chhabra et al., 2022; Moseley et al., 2022, 2014; A R Powers et al., 2017; Schmack et al., 2021). In our two studies with two independent cohorts, our participants detected voices (either their own or someone else’s) presented at individual hearing thresholds in pink noise, while simultaneously experiencing robotic sensorimotor stimulation. We hypothesized that conflicting sensorimotor stimulation leading to a PH (Bernasconi et al., 2022, 2021; Blanke et al., 2014) would lead to an increase in vocal false alarms (i.e., reporting hearing voices in trials with no physical voice present in noise), compared to the stimulation with a weaker sensorimotor conflict, thereby relating our findings with the self-monitoring account. Moreover, we hypothesized that this increase would be modulated by the voice task they are involved in (other-voice detection vs self-voice detection), being especially prominent when performing other-voice detection.

## Method

### Participants

We conducted two studies with the same general procedure and experimental design. Study 2 was set to replicate the effects observed in Study 1. Both studies involved 24 right-handed participants chosen from the general population, fluent in French and naïve to the purpose of the study. In Study 1, 17 participants were female (mean age ± SD: 25.0 ± 4.2 years old), whereas in Study 2, 13 were female (26.6 ± 5.3 years old). Sample size in both studies was similar to our previous work (Orepic et al., 2021) and determined to match the number of all possible permutations of experimental conditions. No participants reported any history of psychiatric or neurological disorders as well as any hearing deficits. Participants gave informed consent in accordance with the institutional guidelines (protocol 2015-00092, approved by the Comité Cantonal d’Ethique de la Recherche of Geneva), and received monetary compensation (CHF 20/h).

### Stimuli

Participants’ voices were recorded (Zoom H6 Handy recorder) while saying nine one-syllable words in French (translated to English: *nail, whip, ax, blade, fight, bone, rat, blood, saw, worm*). The words were chosen from the list of 100 negatively-valenced words, as rated by 20 schizophrenic patients and 97 healthy participants (Jalenques et al., 2013). Negative words were purposefully chosen in our previous study (Orepic et al., 2021), in order to better approximate the phenomenology of AVH, that are mostly negative in content (Woods et al., 2015). After the background noise was removed from the recordings, they were standardized for sound intensity (−12 dBFS) and duration (500 milliseconds) (Audacity software). The preprocessed recordings were used as *self-voice* stimuli in a voice detection task, which also contained *other-voice* stimuli – i.e. equivalent voice recordings of a gender-matched person unknown to the participant. Auditory stimuli were presented to participants through noise-cancelling headphones (Bose QC20). The experimental paradigm was created in MATLAB 2017b with Psychtoolbox library (Brainard, 1997; Kleiner et al., 2007; Pelli, 1997).

### Experimental procedure

Upon arrival, participants were screened for eligibility criteria, after which their voices were recorded. This was followed by two *Sensorimotor blocks* (synchronous and asynchronous), designed to assess illusory effects of sensorimotor stimulation. *Sensorimotor blocks* were followed by *Staircase blocks* (bottom-up and top-down), used to estimate individual hearing thresholds with a voice detection task. Finally, in four *Task blocks* (synchronous-self, synchronous-other, asynchronous-self, asynchronous-other) we assessed vocal false alarms by combining sensorimotor stimulation and voice detection task. At the end of the experiment, participants filled out the PDI questionnaire (Peters et al., 2004), that assesses delusional ideation present in the general population and has been related both to errors in self-monitoring (Teufel et al., 2010) as well as excessive prior-weighting.

### Sensorimotor blocks: assessment of illusory effects

Identical to our previous studies (Blanke et al., 2014; Faivre et al., 2020; Orepic et al., 2021; Salomon et al., 2020; Serino et al., 2021), during sensorimotor blocks participants manipulated a robotic system that consists of two integrated units: the front part – a commercial haptic interface (Phantom Omni, SensAble Technologies) – and the back part – a three degree-of-freedom robot (Hara et al., 2011) (Figure 1). Blindfolded participants were seated between the front and back parts of the robot and were asked to perform repeated poking movements with their right index finger using the front part. Participants’ pokes were replicated by the back part, thus applying corresponding touches on participants’ backs. The touches were mediated by the robot either in synchronous (without delay) or in asynchronous (with 500 milliseconds delay) fashion, creating different degrees of sensorimotor conflict between the upper limb movement and somatosensory feedback on the back.

**Figure 1.**
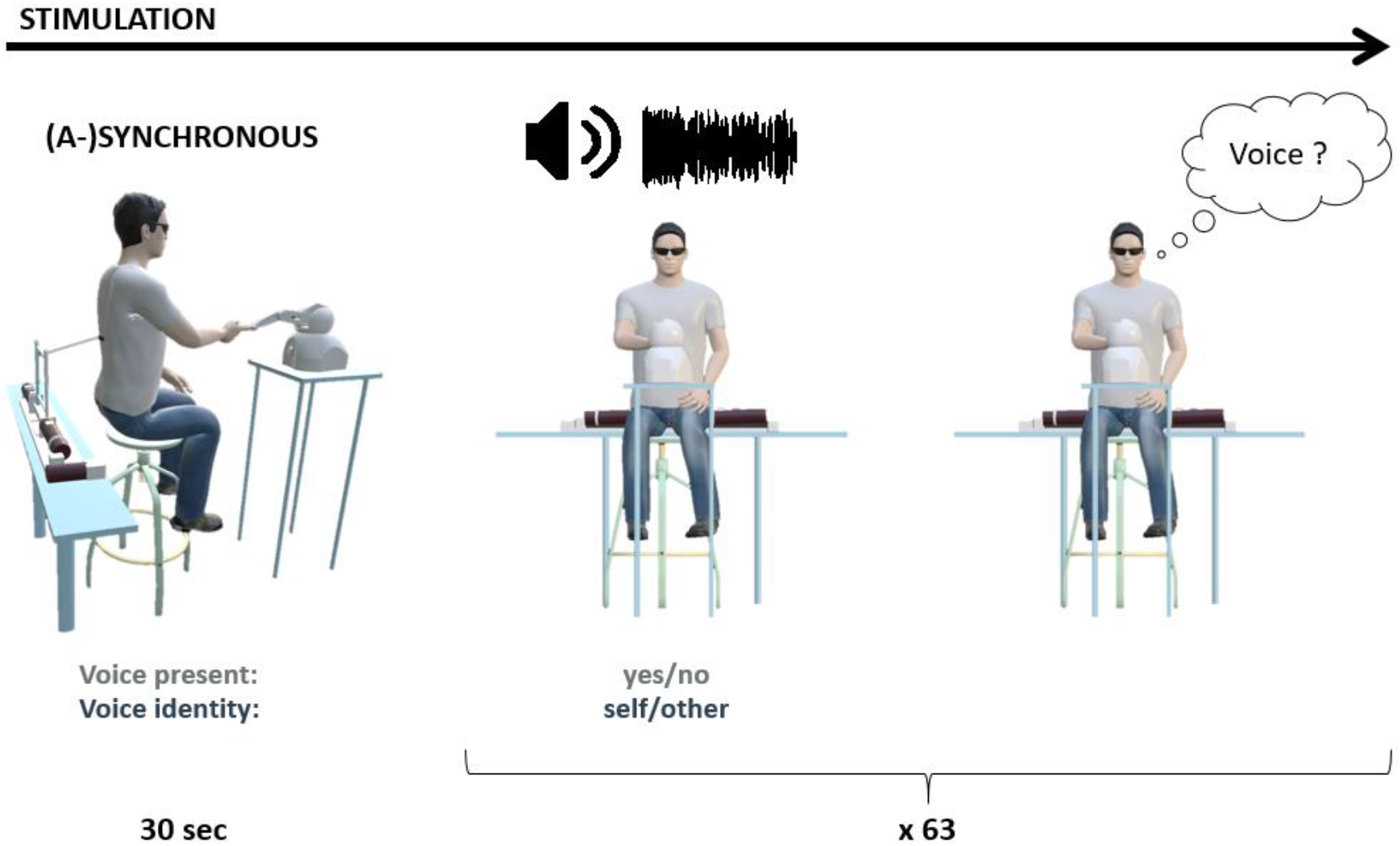
Task block design. The block started with 30 seconds of sensorimotor stimulation, which was followed by simultaneous voice detection task. While manipulating the robotic device, participants were hearing bursts of pink noise and were instructed to report whether they heard a voice in the noise. Out of 63 trials, 45 contained a voice presented at hearing threshold. Within a block, the voices either belonged to participant (self) or to a stranger (other).

Following a two-minute-long sensorimotor stimulation (both synchronous and asynchronous), participants filled out a short questionnaire. Specifically, on a Likert scale from 0 (not at all) to 6 (very strong), after each block, participants rated the strength of illusory self-touch (“I felt as if I was touching my back by myself”), somatic passivity (“I felt as if someone else was touching my back”) and presence hallucination (“I felt as if someone was standing close to me”). Questionnaire contained an additional control item (“I felt as if I had three bodies.”). The order of the two blocks (synchronous and asynchronous) was counterbalanced across participants.

### Staircase blocks: defining hearing thresholds

Participants’ individual hearing thresholds were estimated with a voice detection task combined with a one-up-one-down staircase procedure (Cornsweet, 1962). During the task, participants were continuously hearing short bursts of pink noise and were instructed to report whether they heard a voice in the noise by pressing on a button after the noise offset. Each burst of noise lasted for 3.5 seconds and voice onset randomly occurred in a period between 0.5 and 2.5 seconds after the noise onset, ensuring a minimum of 0.5 seconds of noise before and after the presentation of a voice recording. Following participants’ response in each trial (i.e., a button click after the noise offset), an inter-trial interval jittered between 1 and 1.5 seconds.

The staircase procedure employed only other-voice stimuli and consisted of two blocks, one starting from a high (top-down block) and another from a low (bottom-up block) sound intensity level, counterbalanced across participants. In both staircase blocks, each word was presented four times in a randomized order, resulting in 36 trials. Threshold in each block was computed as a mean value from the last 15 trials and the average of the two thresholds was considered as participants’ hearing threshold. No differences in detectability between self-voice and other-voice stimuli, as well as between different words were assured in a pilot study.

### Task blocks: combining a voice detection task with sensorimotor stimulation

During *Task blocks*, participants were performing the voice detection task while being exposed to sensorimotor stimulation (i.e., while they manipulated the robotic device). *Task blocks* differed based on the type of sensorimotor stimulation (synchronous, asynchronous), as well as of vocal stimuli (self, other). Thus, each participant completed four *Task blocks* (synchronous-self, synchronous-other, asynchronous-self, asynchronous-other) and had a unique order of blocks (i.e., we tested 24 participants to match 24 possible permutations of *Task blocks). Task blocks* started with 30 seconds of sensorimotor stimulation, followed by a concomitant voice detection task (Figure 1). Throughout the auditory task, participants continued manipulating the robot and auditory stimuli were not time-locked to participants’ movements. The voice detection task was identical to the task in *Staircase blocks*, with the addition of 18 trials that contained only noise (i.e., no-voice trials). No-voice trials were randomized together with 45 trials containing a voice (i.e., each word was presented five times within a block), resulting in 63 trials per block. An adaptive one-up-one-down staircase procedure was maintained throughout the block to ensure that the voices were presented at hearing threshold.

### Statistical analysis

Statistical analysis and plotting were performed in R (R Core Team, 2020), using notably the lme4 (Bates et al., 2015), lmerTest (Kuznetsova et al., 2018), and afex (Singmann et al., 2019) packages. The results were illustrated using sjplot (Lüdecke, 2018) and ggplot2 (Wickham, 2016) packages.

### Vocal false alarms

Serving as a measure of experimentally-induced AVH, our primary research interest was to identify the effects of sensorimotor stimulation on vocal false alarm rate. Thus, on no-voice trials, we conducted a mixed-effects binomial regression with Response as dependent variable and Stimulation (synchronous, asynchronous), Voice (self, other) and Gender (male, female) as fixed effects, and participants as random effect. Voice and Stimulation were constant throughout the block. The Response-variable indicates whether participants detected or not a voice in the noise, thus in the no-voice trials it represents a false alarm (whereas for the trials containing a voice in the noise, it stands for a hit). An interaction term was added between the effects of Stimulation and Voice. The Gender effect was added to the regression because of previous reports of gender differences with respect to felt presences as well as AVH (Alderson-Day et al., 2022, 2021). Random effects included a by-participant random intercept. By-participant random slopes for the main effects were added following model selection based on maximum likelihood.

### Questionnaire ratings

Ratings in questionnaire items were assessed by a mixed-effects linear regression containing a fixed effect of Stimulation (synchronous, asynchronous) and by-subject random intercepts. For the questionnaire items that significantly differed between the two sensorimotor stimulations (synchronous, asynchronous), we additionally explored whether the illusion assessed by the corresponding questionnaire item affected false alarm rate in the voice detection task. Specifically, to the mixed-effect binomial regression described above (with Response as a dependent variable) we added an additional fixed effect Questionnaire Item, with values represented as Likert-scale ratings (0-6) given for the corresponding questionnaire item and sensorimotor stimulation. The effect of Questionnaire Item was related with an interaction term with the effect of Condition.

### Delusional ideation

Similar to questionnaire items, we explored the effects of delusional ideation on false alarm rate, by adding PDI score (Peters et al., 2004) as a covariate to the equivalent mixed-effect binomial regression, and forming a two-way interaction together with the effect of Condition.

### Control analyses

Our primary outcome variable was false-alarm rate. However, in order ensure that our experimental manipulation only affected no-voice trials (i.e., false alarms), we also conducted equivalent mixed-effects binomial regression analyses for the trials with voices present in noise (i.e., with hit rate as dependent variable). A lack of equivalent effects on hit rate would indicate that our experimental manipulation did not affect the detectability of the voices when they are actually present in noise, but that the effects are specific to reporting hearing non-existing voices in noise. Besides hit rate, our control variables were d’, i.e., task sensitivity (the distance between the midpoints of distributions of signal and signal with added noise; calculated as the standardized false-alarm rate subtracted from the standardized hit rate) and criterion, i.e., response bias (the number of standard deviations from the midpoint between these two distributions; calculated as the mean of the standardized hit rate and standardized false alarm rate). D’ and criterion were assessed with a two-way ANOVA containing Stimulation (synchronous, asynchronous) and Voice (self, other) as fixed effects with an interaction term.

## Results

### Vocal false alarms

In both studies, we investigated the effects of sensorimotor stimulation (synchronous, asynchronous) and type of voice stimuli (self, other) on the rate of induced vocal false alarms in the voice detection task.

### Study 1

In Study 1, a mixed-effects binomial regression revealed a main effect of Stimulation (estimate = −0.58, Z=−2.12, p=0.034), indicating a higher false alarm rate during asynchronous (mean = 0.15, 95% CI = [0.13, 0.17]), compared to synchronous stimulation (mean = 0.13, 95% CI = [0.11, 0.16]). Critically, there was a significant interaction between Voice and Stimulation (estimate=1.09, Z=2.72, p=0.007) (Figure 2A). Post-hoc analyses of the interaction indicated that during the blocks containing other-voice stimuli, false alarm rate was higher with asynchronous stimulation (estimate=−0.56, Z=−2.1, p=0.036; mean = 0.16, 95% CI = [0.13, 0.19], synchronous: mean = 0.12, 95% CI = [0.09, 0.15]). During self-voice blocks, there was a tendency for the opposite effect: an increase in false alarm rate with synchronous stimulation (estimate=0.52, Z=1.77, p=0.077; mean = 0.16, 95% CI = [0.13, 0.19], asynchronous: mean = 0.14, 95% CI = [0.11, 0.17]) (Figure 2A). There was no significant main effect of Voice (estimate=−0.87, Z=−1.83, p=0.068). This shows that 1) there were more vocal false alarms during the condition with higher sensorimotor conflict, and 2) this was more prominent in the blocks containing other-voice stimuli.

**Figure 2.**
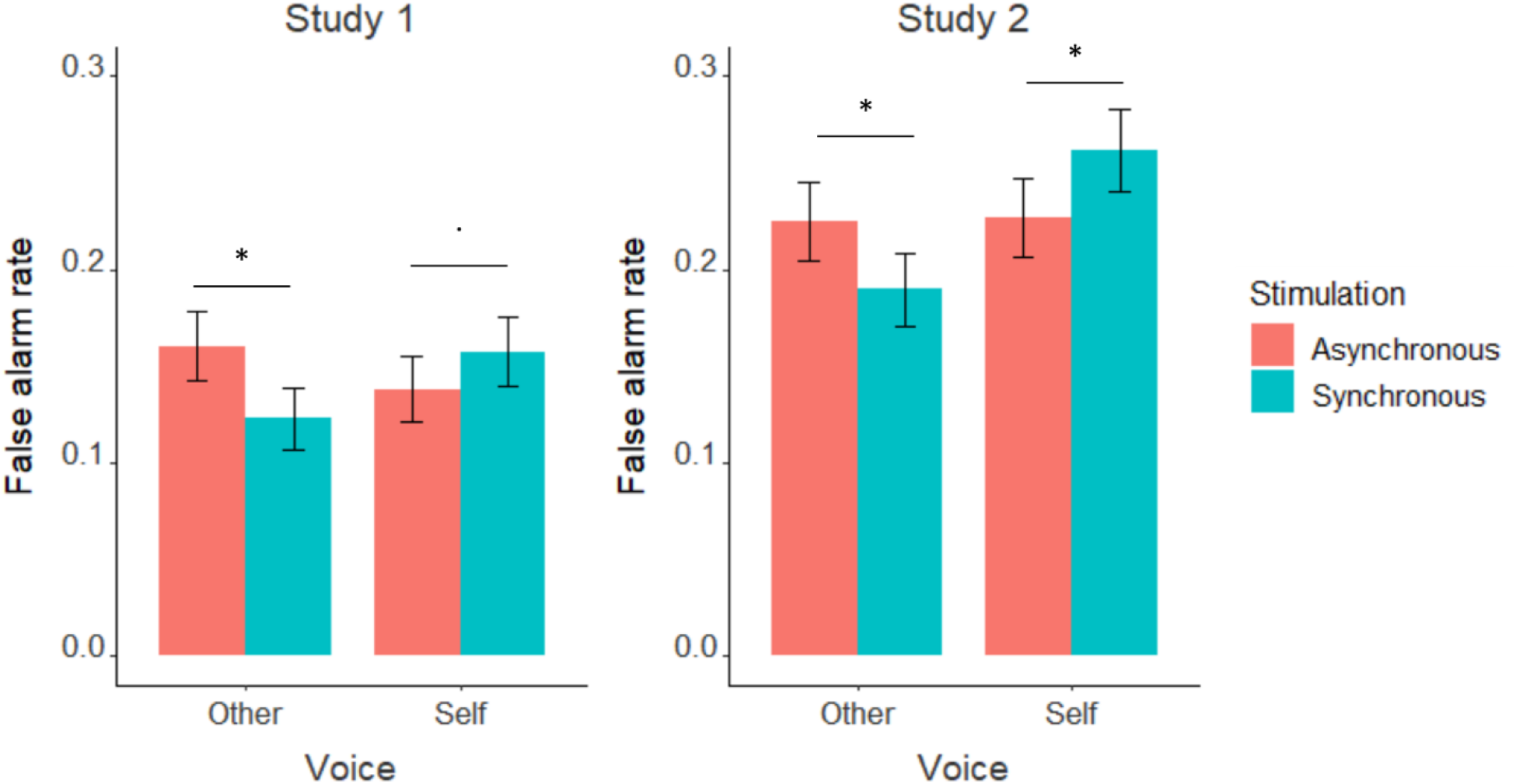
Vocal false alarm rates observed in Study 1 (left) and Study 2 (right). Height of bar plots indicates mean rate, and error bars 95% confidence intervals. In both studies, asynchronous stimulation increased false alarm rate in blocks containing other-voice stimuli, whereas synchronous stimulation increased false alarms in self-voice blocks. *: p < 0.05,.: p < 0.1

### Study 2

In Study 2, we replicated this interaction effect between Stimulation and Voice (estimate=1.14, Z=2.97, p=0.003), again revealing that in other-voice blocks false alarms increased with asynchronous (estimate=−0.64, Z=−2.15, p=0.031, mean = 0.22, 95% CI = [0.19, 0.25], synchronous: mean = 0.19, 95% CI = [0.16, 0.22]), whereas in self-voice blocks the opposite effect occurred – more false alarms in synchronous stimulation (estimate=0.53, Z=1.98, p=0.048; mean = 0.26, 95% CI = [0.22, 0.3], asynchronous: mean = 0.23, 95% CI = [0.19, 0.27]) (Figure 2B). Also, there were, again, more false alarms during asynchronous stimulation (main effect of Stimulation; estimate=−0.63, Z=−2.21, p=0.027; mean = 0.23, 95% CI = [0.2, 0.23], synchronous: mean = 0.23, 95% CI = [0.2, 0.23]). There were no differences in false alarms between the two voices (estimate=0.03, Z=0.14, p=0.887). Neither study had a significant effect of Gender (all p > 0.05, Supplementary material). This shows that the same effects of sensorimotor stimulation on vocal false alarms were replicated in an independent cohort of participants.

### Delusional ideation

We also investigated the potential relationship between the observed increase in vocal false alarms and delusional ideation (measured by the PDI questionnaire (Peters et al., 2004)). In both independent subject samples that we tested (Study 1 and 2), binomial mixed-effect regression of responses in non-voice trials revealed a significant main effect of PDI score (Study 1: estimate=0.2, Z=2.17, p=0.03; Study 2: estimate=0.33, Z=2.15, p=0.032), indicating that the higher participants scored on delusional ideation inventory, the more false alarms they made during the voice detection task. In both studies, there was a tendency for a significant interaction between the effects of Stimulation and PDI (Study 1: estimate=−0.2, Z=−1.9, p=0.057; Study 2: estimate=−0.1, Z=−1.9, p=0.058), indicating that this relationship was stronger during asynchronous stimulation (Figure 3). We also divided the PDI scores into 3 subcategories – distress, preoccupation, and conviction (Peters et al., 2004; Schmack et al., 2013) – and ran equivalent mixed effect analyses. The only consistent result across the two studies was an interaction between Stimulation and Conviction score (Study 1: estimate=−0.08, Z=−2.53, p=0.011; Study 2: estimate=−0.04, Z=−2.17, p=0.029), indicating a stronger relationship between Conviction and false alarms during Asynchronous stimulation. The results for other subcategories are reported in Supplementary material.

**Figure 3.**
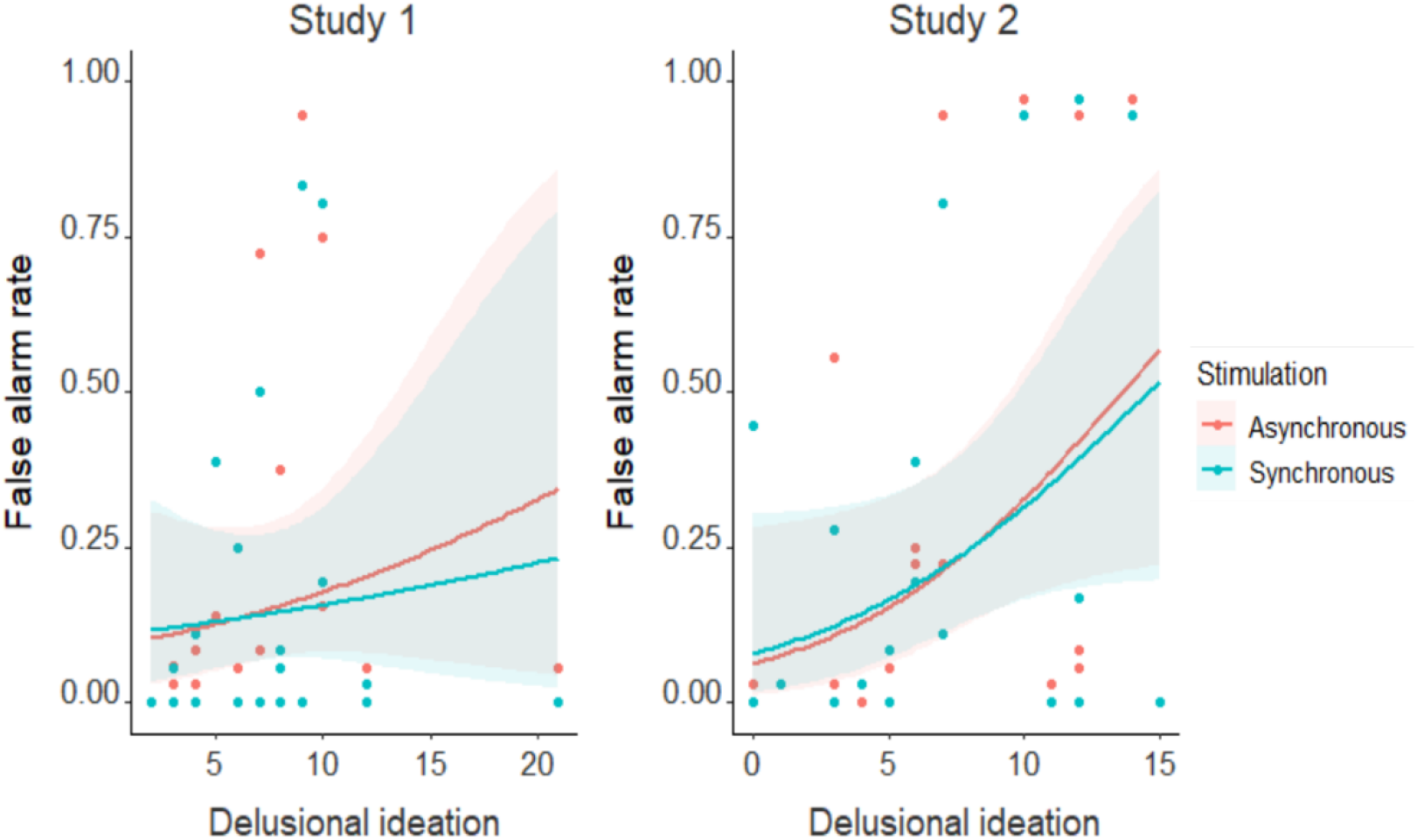
Increase in delusional ideation score was related to an increase in vocal false alarms rate in both studies. Shaded areas around each curve represent 95% confidence intervals.

Together, these data show that increased false alarms are related to delusional ideation, suggesting a presence of perceptual priors underlying the effects on robotically-induced false alarms.

### Questionnaire ratings

Presence hallucination was, as expected, experienced more during asynchronous stimulation (Study 1: estimate=−0.5, t(24)=−2.68, p=0.013; Study 2, tendency: estimate=−0.67, t(24)=−1.92, p=0.067). Somatic passivity was also rated higher during asynchronous stimulation (Study 1: estimate=−1.08, t(24)=−3.68, p=0.001; Study 2 tendency: estimate=−0.58, t(24)=−1.81, p=0.083), whereas Self-touch ratings were higher during synchronous compared to asynchronous stimulation (Study 1: estimate=0.79, t(24)=2.21, p=0.037; Study 2: estimate=0.83, t(24)=3.46, p=0.002). The control questionnaire item was unaffected by sensorimotor stimulation (Study 1: estimate=−0.04, t(24)=−0.58, p=0.566; Study 2: estimate=0.13, t(24)=1.39, p=0.176). There were no significant effects of gender (all p > 0.05). Means and standard deviations of all questions for both studies are reported in Supplementary material. Correlation between questionnaire ratings and false alarm rates indicated results that were inconsistent between the two studies and are thus reported in Supplementary material. Together, this shows that in both studies, our robotic stimulation induced the expected experiential consequences (Bernasconi et al., 2021; Blanke et al., 2014) – i.e., higher PH and somatic passivity with asynchronous stimulation.

### Control analyses

As control analyses, we investigated the effects on Stimulation and Voice on hit rate (i.e., responses in trials where there was voice present in noise), as well as on d’ and criterion. In Study 1, binomial mixed-effects regressions of responses in voice-trials revealed a tendency for a main effect of Stimulation (estimate=−0.23, Z=−1.94, p=0.052), underlying a higher hit rate during asynchronous (mean = 0.55, 95% CI = [0.53, 0.57]) than synchronous stimulation (mean = 0.53, 95% CI = [0.51, 0.55]). Hit rates were unaffected by Voice (estimate=0, Z=0.03, p=0.979) and Gender (estimate=−0.03, Z=−0.11, p=0.91). The interaction between Voice and Stimulation indicated a tendency towards significance (estimate=0.23, Z=1.81, p=0.071). In Study 2, none of the tendencies from Study 1 proved significant. There was no main effect of Stimulation (estimate=−0.08, Z=−0.76, p=0.446), nor it interacted with Voice (estimate=0.12, Z=0.9, p=0.367). There were no significant effects of Gender (estimate=−0.7, Z=−1.73, p=0.084) nor Voice (estimate=−0.09, Z=−0.81, p=0.416).

In neither study did we observe significant effects on d’ and criterion (Supplementary material).

The control analyses show that the detectability of the voices, when they are indeed present in noise (i.e., hit rate), is affected neither by Stimulation nor type of detected voice, and that the aforementioned effects are specific to trials in which there is no voice present in noise (i.e., on false alarms).

## Discussion

In two independent cohorts of healthy participants, we employed specific sensorimotor robotic stimulation to induce AVH-like phenomena, as indicated by an increase in the false alarm rate while participants were engaged in a voice detection task. Critically, the presence of AVH-like phenomena was additionally modulated by the type of sensorimotor stimulation. Thus, the asynchronous sensorimotor condition (related to other-agent sensations; PH and somatic passivity) induced overall more AVH-like phenomena and this effect was especially observed in experimental blocks containing other-voice stimuli. Finally, the rate of AVH-like phenomena was positively related to delusional ideation and this correlation was stronger for vocal false alarms during the asynchronous PH-inducing condition.

Extending our sensorimotor procedure that has been shown to induce PH in healthy subjects (Bernasconi et al., 2022; Blanke et al., 2014; Dhanis et al., 2022; Orepic et al., 2021; Serino et al., 2021) and patients with Parkinson’s disease (Bernasconi et al., 2021), we here demonstrate a new experimental paradigm able to induce controlled AVH-like phenomena (manifested as specific false alarms) in healthy, non-hallucinating individuals. Previous methods of inducing hallucinations in healthy individuals – such as through psychedelic medications (Preller and Vollenweider, 2018) or by automatized visual stimulations (e.g., Flicker-induced (Allefeld et al., 2011; Pearson et al., 2016)) – have identified many important challenges present in hallucination engineering. These are, for instance, difficulties to repeatedly induce a hallucination within a given and short period of time, investigating the hallucination of a given participant in real time, and quantifying hallucinations with objective measures (as opposed to measures such as verbal self-reports) – rendering them prone to participant and experimenter biases (Adler, 1973; Rosenthal and Fode, 1963). Our AVH-inducing paradigm partly address these challenges – e.g., we were able to repeatedly elicit a perception of non-existing voices in short experimental blocks, which were quantified objectively (through a false alarm rate) and in real time. Moreover, we elicited hallucinations in healthy participants, thereby controlling for confounds related to disease, in which hallucinations typically occur. Finally, our approach on AVH-like experiences allowed us to compare our main findings with the two most prominent views about AVH – the self-monitoring account and the strong-priors account, providing evidence for the clinical relevance of both theoretical accounts.

According to the self-monitoring account, AVH result from aberrant predictive mechanisms related to motor actions and the corresponding sensory feedback (Miall and Wolpert, 1996; Shadmehr et al., 2010; Wolpert et al., 1995). Evidence from previous studies arguing for the self-monitoring account is largely implicit, and mainly links AVH to either 1) differences in the amplitude of auditory evoked responses between self- and externally-generated actions or 2) differences in interregional communication within the speech network (Ford et al., 2012, 2002; Friston and Frith, 1995; Hoffman et al., 2011; Whitford, 2019). Here, however, we explicitly manipulate self-monitoring with our robotic procedure, and, as a consequence, observe differences in the magnitude of experimentally-evoked AVH-like sensations (i.e., vocal false alarms). Specifically, with our robotic procedure, we induced two levels of self-monitoring perturbation – with stronger (asynchronous) and weaker (synchronous) sensorimotor conflicts – and observed a differential effect on AVH-like sensations (i.e., more false alarms with stronger sensorimotor conflicts). This shows that healthy individuals are more likely to hear non-existing voices in auditory noise when simultaneously experiencing self-monitoring perturbations, which suggests that similar mechanisms might occur in the brain of voice hearers. It is important to note, however, that AVH as quantified in the present experiments arose not as an impairment in speech-related sensory predictions in voice perception, but as an alteration of tactile, proprioceptive, and motor processes involved in self-monitoring (related to the repetitive arm movements and their corresponding tactile feedback on the back), which was sufficient to alter voice perception, as previously shown for the clinically related phenomenon of thought insertion (Serino et al., 2021). This extends the current view of the self-monitoring account for AVH to a more global deficit in self-monitoring (Blanke, 2012; Blanke et al., 2015; Serino et al., 2021), that has until now been almost exclusively focused on predictions related to speech-related signals (Ford and Mathalon, 2019; Whitford, 2019). Collectively, these findings suggest that errors in self-monitoring, induced by conflicting sensorimotor stimulations involving arm and trunk signals, and even if not involving an experimental manipulation of auditory-verbal signals, are sufficient to exert a specific effect on auditory-verbal perception, reflected as a proneness to hear non-existing voices in noise.

Concerning the strong-priors account, AVH are proposed to arise when strong beliefs (i.e., priors) exert a top-down effect on perception (Cassidy et al., 2018; Corlett et al., 2019; A R Powers et al., 2017; Zarkali et al., 2019). Evidence from previous studies arguing for the strong-priors account mainly links AVH to favoring prior knowledge over sensory information during perceptual inference (Alderson-Day et al., 2017; Kot and Serper, 2002; Teufel et al., 2015). In a classical experimental scenario, researchers experimentally manipulate trial-by-trial multisensory (e.g., visuo-auditory) contingencies (Davies et al., 1982; Ellson, 1941; A R Powers et al., 2017) to engender stronger priors about the occurrence of the hallucinated auditory stimulus (e.g., a false alarm in a tone-detection task). However, these studies did not investigate which type of a perceptual prior can lead to which type of a hallucinated stimulus, especially with respect to vocal stimuli. Here, we extend this work by showing that different auditory-perceptual priors induced through our robotic stimulation can affect the type of AVH-like sensations (i.e., self-voice false alarm versus other-voice false alarm). Specifically, during asynchronous stimulation, that is related to otherness-related sensations (i.e., PH and somatic passivity) (Bernasconi et al., 2021; Blanke et al., 2014; Serino et al., 2021), there were more false alarms in other-voice blocks, compared to self-voice blocks. Thus, while exposed to asynchronous PH-inducing stimulation, participants reported hearing a non-existing voice in noise more often in blocks where noise stimuli (without any voice) were mixed with other-voice stimuli, compared to self-voice stimuli. Accordingly, we propose that our robotic procedure induces a perceptual prior (otherness sensations) that, in turn, exerts a specific top-down effect on auditory-verbal perception leading to false alarms of another voice. As a control, during synchronous stimulation, that is associated to self-touch sensations, we observed more false alarms in self-voice blocks.

AVH-like sensations induced through our paradigm might relate the self-monitoring and strong-priors accounts. Namely, in our paradigm, there might be two perceptual priors at play – an auditory-verbal prior and a sensorimotor prior (coming from the robotic stimulation). The auditory-verbal prior consists of the fact that repeatedly hearing a voice with a specific identity (self or other) throughout our experimental blocks creates an expectation about the identity of the voices to follow – i.e., if one continuously hears consecutive other-voice stimuli, one might expect to hear other-voice again in the near future. Concerning the sensorimotor prior, it has been proposed that there might be different intersecting hierarchies in the brain – related to self-monitoring, selfrelated priors and other-related priors – and that errors in self-monitoring are explained away by changing precision either of self-related or other-related priors (Corlett et al., 2019; Leptourgos and Corlett, 2020). Applied to our data, self-monitoring perturbations coming from asynchronous stimulation might thus be explained away by increasing precision of other-related priors (e.g., perceived as PH). Crucially, the directionality of the imposed ‘voice prior’ (self or other) might hence interfere with the prior stemming from the concomitant sensorimotor stimulation, moreover in a complementary fashion. Thus, other-voice prior combined with other-related sensorimotor prior might lead to an increase in other-voice false alarms. This proposed mechanism is illustrated on Figure 4.

**Figure 4:**
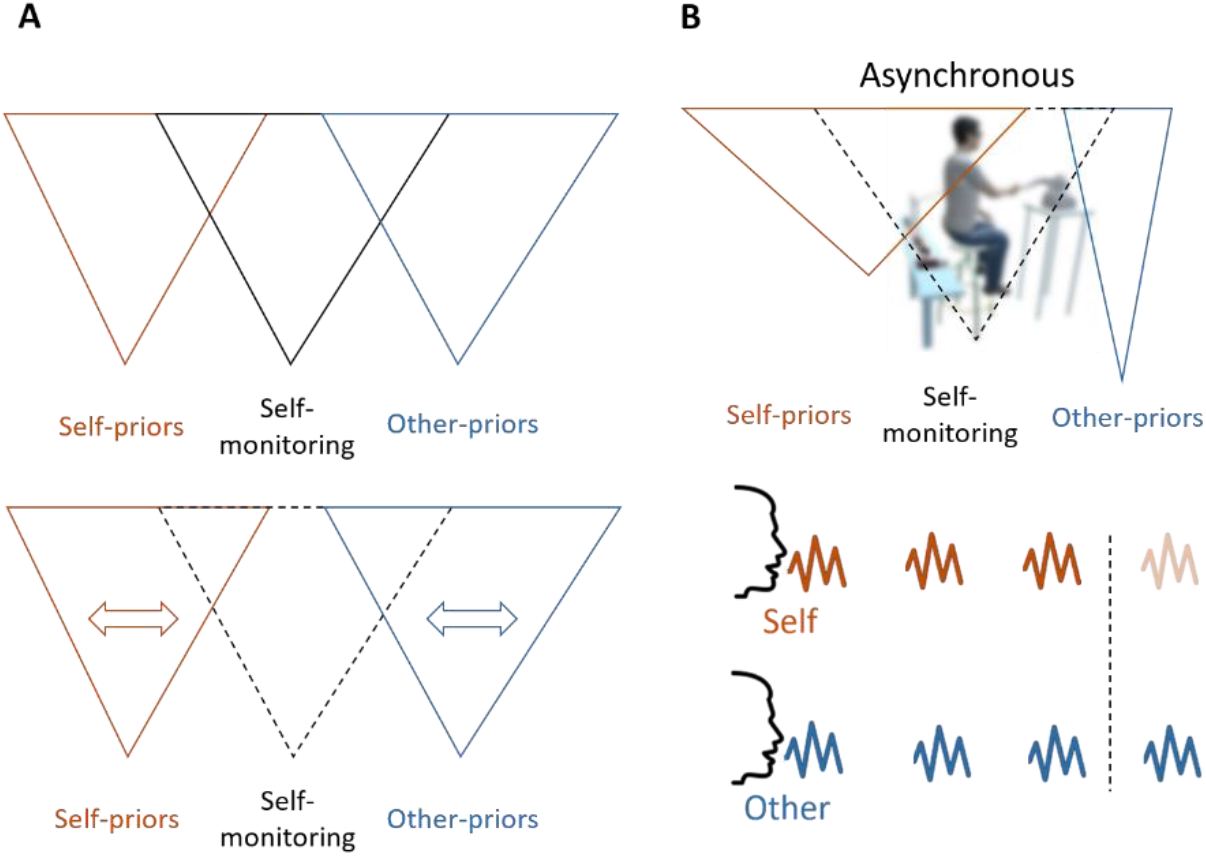
Proposed mechanism for the observed identity-specific vocal false alarms. **A)** Top: The pyramids indicate intersecting hierarchies for processing of self-monitoring, self-priors and other-priors, proposed by (Leptourgos and Corlett, 2020). Bottom: Errors in self-monitoring hierarchy (dashed lines) are explained away by changes in precision of self- and other-priors, resulting in self- or other-attribution biases (changes in the width of the corresponding pyramid). **B)** Top: Self-monitoring errors during asynchronous stimulation are explained away by increasing precision of other-related priors (narrower other-priors pyramid). Bottom: Repeated exposure to the same type of voice (self or other) drives an expectation to hear the same type of voice in the near future (after the vertical line). Concomitant increase in other-priors’ precision imposes an expectation to hear other-voice (blue), as opposed to self-voice (orange), resulting in increased other-voice false alarms (opaque color).

Relationship between robot-induced effects and beliefs is further corroborated in our data by relating the increase of false alarms with PDI score that reflects delusion proneness. Namely, delusional ideation has been related both to endorsement of hallucinations (Varghese et al., 2008) and to excessive prior-weighting (McLean et al., 2020; Schmack et al., 2015; Teufel et al., 2015) in the general population. This suggests that PDI might measure a trait of overly strong perceptual priors that impose top-down effects on perception (Adams et al., 2015; Fletcher & Frith, 2009; Sterzer et al., 2018), in addition to the top-down effect (state) induced by sensorimotor stimulation. Moreover, from the three subscales of delusional ideation – distress, conviction, and preoccupation – a consistent relationship to false alarms similar to the general PDI score was observed only for the conviction scale. Conviction subscale was previously related to a reduced perceptual stability and a stronger belief-induced bias on perception, as well as stronger functional connectivity between frontal areas encoding beliefs and sensory areas encoding perception (Schmack et al., 2013), suggesting that similar effects might be at play here.

Our findings further support previous studies that argue for the use of auditory false alarms as a proxy for hallucinations. In a large-scale multisite study (Moseley et al., 2021), hallucinatory experiences in more than a thousand participants were associated with false alarms in a signal detection task, as opposed to other commonly used hallucination-related tasks (e.g., source memory or dichotic listening). Similarly, in a recent review, signal and voice detection tasks were found to be the most robust amongst five most common voice-hearing induction paradigms (Anderson et al., 2021). Importantly, in our studies, auditory detectability between experimental stimuli (self and other voice) and sensorimotor stimulations (synchronous and asynchronous) cannot account for present effects, as sensorimotor stimulation did not affect hit rate in voice detection task. Specifically, when there were voices present in the noise, participants were able to detect them equally well, regardless of the concomitant sensorimotor stimulation. Differences in sensorimotor stimulation only affected the performance in trials with no voices present in noise.

In conclusion, here we demonstrate a sensorimotor-robotic procedure and method (Bernasconi et al., 2022) that is able to induce AVH-like sensations in healthy individuals and in a fully controlled laboratory environment. Specifically, we show that different types of sensorimotor stimulation can selectively induce vocal false percepts and that stimulations that induce sensations related to otherness and an alien agent led to a higher number of other-voice false alarms, an effect that we further related with delusion proneness. Besides the novelty and the important methodological impact, these results shed new light on AVH phenomenology, providing experimental support for both prominent albeit seemingly opposing accounts – portraying AVH as a hybrid between deficits in self-monitoring and hyper-precise priors.

## Author contribution

Study concept and design: PO, NF, OB. Acquisition of data: PO, MF. Analysis and interpretation of data: PO, FB, OB. Drafting of the manuscript: PO, FB, OB. Critical revision of the manuscript for important intellectual content: All authors. Statistical analysis: PO, FB. Obtained funding: OB. Administrative, technical, or material support: All authors. Study supervision: OB.

