## supplementary material for "Robotically-induced auditory-verbal hallucinations: combining self-monitoring and strong perceptual priors"

Olaf Blanke

Bertarelli Chair in Cognitive Neuroprosthetics, Neuro-X Institute & Brain Mind Institute, School of Life Sciences, Campus Biotech, École Polytechnique Fédérale de Lausanne (EPFL), 1012 Geneva, Switzerland

### Questionnaire ratings

Supplementary Table T1 reports means and standard deviations of all questionnaire ratings for both studies and Supplementary Figure S1 illustrates the ratings.

**Table T1.** Mean and standard deviation (SD) of questionnaire items assessed in sensorimotor blocks in both studies.

| | *Study 1* | | | | *Study 2* | | | |
|---|---|---|---|---|---|---|---|---|
| | Asynchronous | | Synchronous | | Asynchronous | | Synchronous | |
| | Mean | SD | Mean | SD | Mean | SD | Mean | SD |
| *Self-touch* | 2.08 | 2.22 | 2.88 | 2.29 | 1.21 | 1.74 | 2.04 | 2.1 |
| *Somatic passivity* | 3.42 | 2.12 | 2.33 | 2.09 | 3.71 | 2.1 | 3.13 | 2.29 |
| *Presence Hallucination* | 1.75 | 1.92 | 1.25 | 1.7 | 2.54 | 2 | 1.88 | 2.29 |
| *Control* | 0.21 | 0.59 | 0.17 | 0.48 | 0.17 | 0.48 | 0.29 | 0.81 |

In Study 1, binomial mixed-effects analysis investigating the effects of questionnaire ratings on false alarm rates, with Response as dependent variable and Stimulation and Somatic Passivity as fixed effects, indicated a significant interaction between Somatic Passivity and Stimulation (estimate=-0.53, Z=-4.3, p<0.001). This effect, however, was not replicated in Study 2 (estimate=-0.04, Z=-0.35, p=0.724). Similarly, in Study 1, there was a tendency for an interaction between Presence Hallucination and Stimulation (estimate=-0.35, Z=-1.78, p=0.076) that was not replicated in Study 2 (estimate=-0.17, Z=-1.47, p=0.143). These interactions in Study 1 indicated

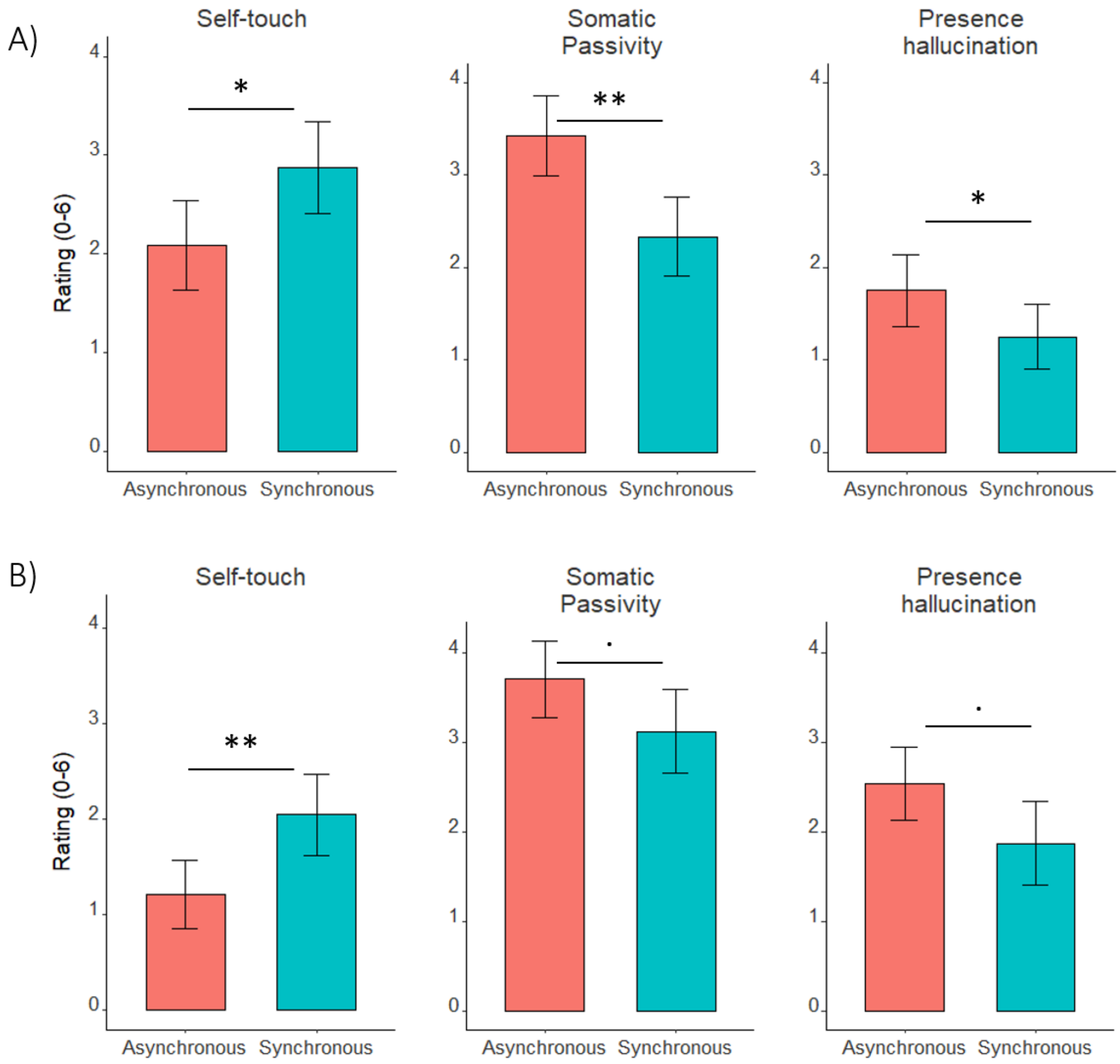


**Figure S1.** Illusory effects assessed in sensorimotor blocks Study 1 (A) and Study 2 (B). Height of bar plots indicates mean rating, and error bars 95% confidence intervals. In both studies, self-touch was higher during synchronous, whereas somatic passivity and presence hallucination during asynchronous stimulation. **: p<0.01, *:p<0.05, .:p<0.1

a stronger relationship between Somatic Passivity and Presence Hallucination with false alarms in Asynchronous, compared to Synchronous stimulation. Self-touch did not have any effects on false alarms in either study (all p > 0.05).

### Signal detection theory

In neither study did we observe significant effects on d' (Supplementary Table T2) and criterion (Supplementary Table T3). However, as in both studies there were tendencies for a main effect of Stimulation only in other-voice trials (Study 1 Self: $F(1, 23)=0.17$, $p=0.685$; Study 1 Other: $F(1, 23)=3.6$, $p=0.071$; Study 2 Self: $F(1, 23)=0.73$, $p=0.403$; Study 2 Other: $F(1, 23)=2.95$, $p=0.099$;), indicating a lower criterion for asynchronous compared to synchronous stimulation (Supplementary Figure S2), we additionally ran exploratory analysis of d' and criterion by merging the data from both studies. This analysis revealed a significant interaction between Stimulation and Voice for the criterion value ($F(1, 47)=4.11$, $p=0.048$). Further investigation of this interaction revealed that criterion was reduced in the asynchronous (mean = 0.44, 95% CI = [0.19, 0.69]) compared to synchronous condition (mean = 0.57, 95% CI = [0.35, 0.79]) only for other-voice blocks (F(1, 47 =6.36, $p=0.015$), with no differences between the two stimulations for self-voice blocks ($F(1, 47)=0.79$, $p=0.38$; asynchronous: mean = 0.50, 95% CI = [0.25, 0.75]; synchronous: mean = 0.43, 95% CI = [0.16, 0.70]). There were no significant main effects of Stimulation ($F(1, 47)=0.35$, $p=0.558$) and Voice ($F(1, 47)=0.81$, $p=0.372$) on the criterion value with merged studies, as well as no significant effects on d' (Condition: ($F(1, 47)=0.09$, $p=0.761$; Voice: ($F(1, 47)=0.23$, $p=0.632$; Condition * Voice: ($F(1, 47)=1.18$, $p=0.283$).

Lower criterion in asynchronous condition suggests a liberal observer, i.e. proneness to respond 'yes'. Thus, these results suggest that in other-voice blocks, participants were more likely to report hearing a voice during asynchronous, compared to synchronous stimulation Although this analysis is exploratory and indicates weak results, these findings are in accordance with the results reported in the main text.

**Table T2.** A two-way ANOVA assessing d’ in both studies.

| | *Study 1* | | | *Study 2* | | |
|---|---|---|---|---|---|---|
| | df | F | p | df | F | p |
| *Stimulation* | 1, 23 | 0.1 | 0.759 | 1, 23 | 0 | 0.95 |
| *Voice* | 1, 23 | 1.27 | 0.272 | 1, 23 | 3.97 | 0.058 |
| *Stimulation * Voice* | 1, 23 | 0.2 | 0.66 | 1, 23 | 1.45 | 0.241 |

**Table T3.** A two-way ANOVA assessing criterion in both studies.

| | *Study 1* | | | *Study 2* | | |
|---|---|---|---|---|---|---|
| | df | F | p | df | F | p |
| *Stimulation* | 1, 23 | 0.1 | 0.759 | 1, 23 | 0 | 0.95 |
| *Voice* | 1, 23 | 1.27 | 0.272 | 1, 23 | 3.97 | 0.058 |
| *Stimulation * Voice* | 1, 23 | 0.2 | 0.66 | 1, 23 | 1.45 | 0.241 |

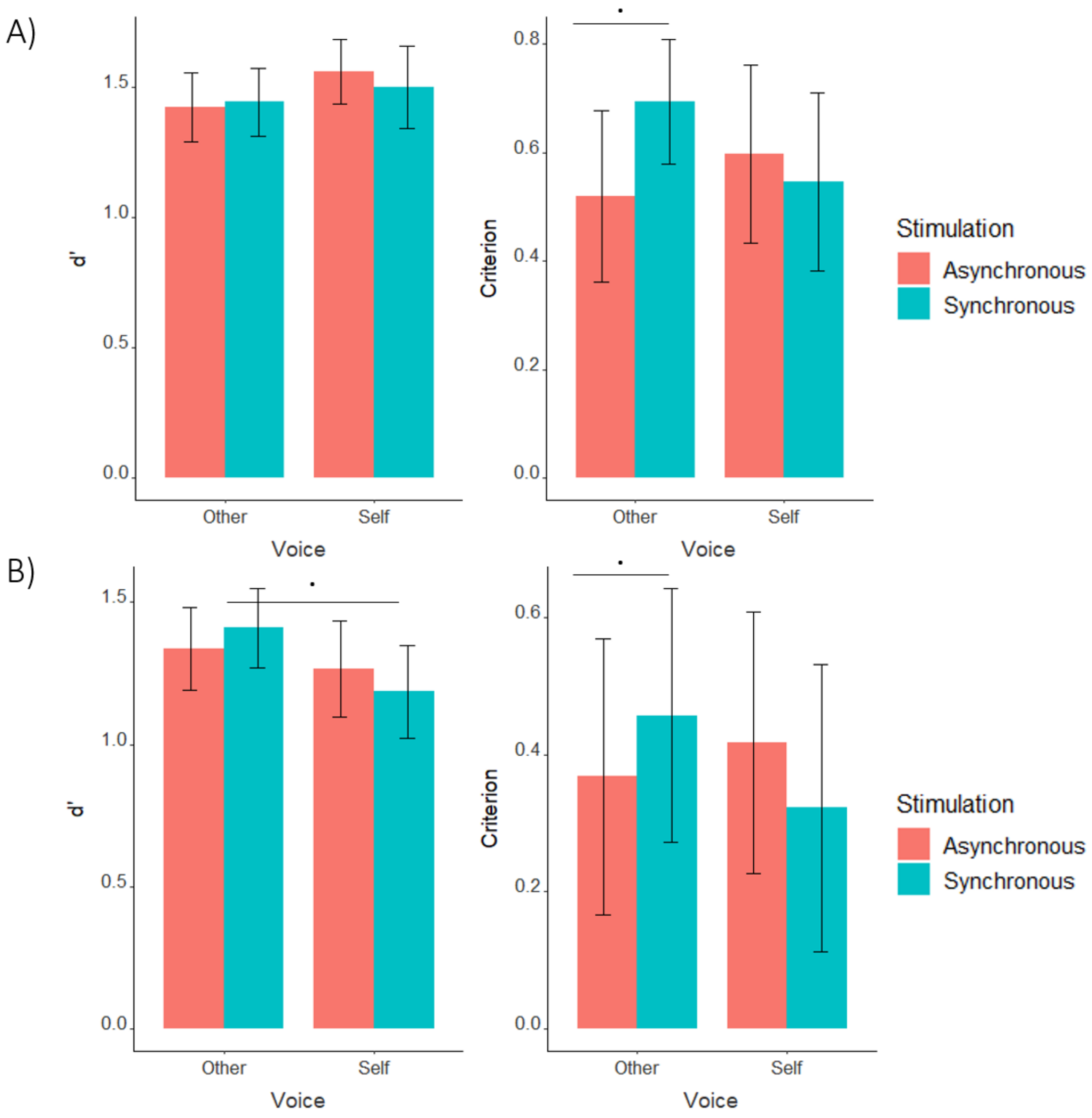


**Figure S2.** Criterion and d' values in both Study 1 (A) and Study 2 (B). In both studies, there were tendencies for a reduced criterion in Asynchronous compared to Synchronous stimulation, but only in Other-voice blocks. In Study 2, there was a tendency for a reduced d' in Self-voice compared to Other-voice blocks. .:p<0.1

#### PDI subcategories

Besides the general PDI score, PDI questionnaire contains three subcategories – distress, preoccupation and conviction (Peters et al., 2004). Previous work has indicated effects of beliefs on perception that were associated specifically with the conviction subscore, in addition to the general PDI score (Schmack et al., 2013). Similarly, we found a significant interaction between the fixed effects Conviction and Stimulation – indicating a stronger positive relationship between false alarm rate and conviction score in asynchronous, compared to synchronous stimulation (Supplementary Figure S3, details in the main text). In Study 1, we observed equivalent interactions with the effect of Stimulation for both effects of Distress (estimate=-0.1, Z=-2.77, p=0.006) and Preoccupation (estimate=-0.13, Z=-3.61, p<0.001), again indicating a stronger relationship between PDI sub-scores and false alarms during Asynchronous stimulation. These interactions were, however, not replicated in Study 2 (distress: estimate=-0.03, Z=-0.99, p=0.325; preoccupation: estimate=-0.03, Z=-1.64, p=0.1). Study 1 also indicated tendencies for the main effects of all 3 PDI subcategories (distress: estimate=0.08, Z=1.85, p=0.065; preoccupation: estimate=0.08, Z=1.92, p=0.054; conviction: estimate=0.07, Z=1.65, p=0.099), indicating a general increase of false alarms with the increase of the scores. In Study 2, only the main effect of distress proved to be significant (distress: estimate=0.11, Z=2.35, p=0.019; preoccupation: estimate=0.08, Z=1.52, p=0.129; conviction: estimate=0.04, Z=0.85, p=0.395).

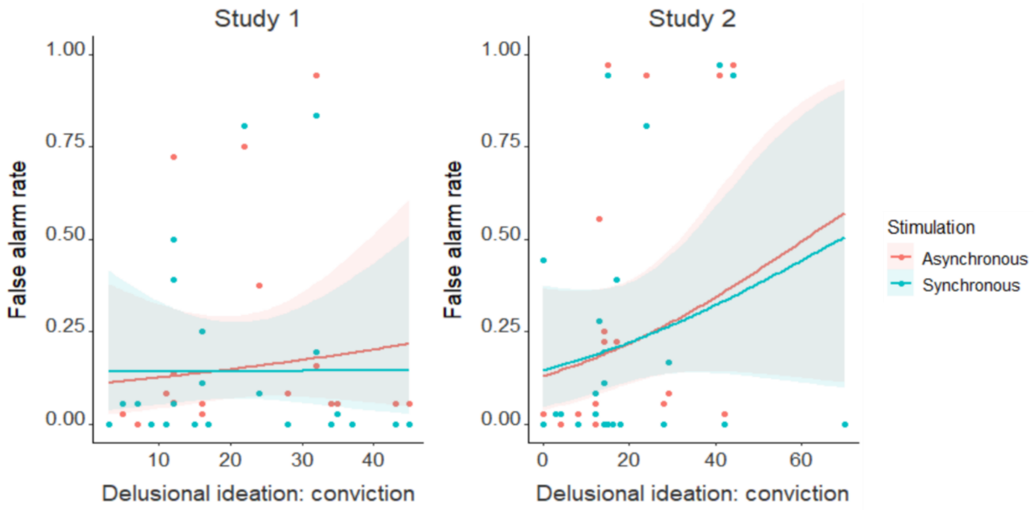


**Figure S3**. In both studies, increase in PDI conviction score was more strongly related to an increase in vocal false alarms rate during asynchronous, compared to synchronous stimulation. Shaded areas around each curve represent 95% confidence intervals.

### Gender effects

Previous work has indicated that females are more likely to experience PH (Alderson-Day et al., 2022) and personified voices (Alderson-Day et al., 2021), compared to males. In Study 1, we found a tendency for the corresponding effect of Gender – false alarms were more often reported by female participants (estimate=-2.05, Z=-1.88, p=0.06). However, this effect was not replicated in Study 2 (estimate=-1, Z=-0.71, p=0.479), and could have arisen from the fact that in Study 1 there were more female participants. When performing exploratory analysis by pooling the participants from both studies together, the effect of gender was (strongly) significant (see below). There were no significant effects of gender on questionnaire ratings.

#### Merging both studies

Since experimental design was equivalent in both studies, as exploratory analysis, we analyzed both studies together – i.e. we placed participants from both studies in the same mixed-effects binomial regression. The effects reported in the main text were more significant compared to having separate models for each study (there were more FAs during asynchronous stimulation and this was more prominent in other-voice blocks), and the gender effect was significant (there were more false alarms in female participants). The model is summarized in Supplementary table T4. Post-hoc analysis of the interaction between Stimulation and Voice showed a significant effect of Stimulation in other-voice blocks (estimate=-0.33, Z=-2.25, p=0.024), again indicating more false alarms during asynchronous stimulation. In self-voice blocks, there was a tendency for the opposite effect – more false alarms during synchronous stimulation (estimate=0.25, Z=1.78, p=0.075).

**Table T4.** Binomial mixed-effects regression for merged studies 1 and 2.

| | *estimate* | *Z* | *p* |
|---|---|---|---|
| *Stimulation* | -0.32 | -2.22 | 0.026 |
| *Voice* | -0.19 | -0.85 | 0.394 |
| *Stimulation * Voice* | 0.57 | 2.82 | 0.005 |
| *Gender* | -0.82 | -5.36 | < 0.001 |
